# A neural mechanism for terminating decisions

**DOI:** 10.1101/2022.05.02.490327

**Authors:** Gabriel M Stine, Eric M Trautmann, Danique Jeurissen, Michael N Shadlen

## Abstract

The brain makes decisions by accumulating evidence until there is enough to stop and choose. Neural mechanisms of evidence accumulation are well established in association cortex, but the site and mechanism of termination is unknown. Here, we elucidate a mechanism for termination by neurons in the primate superior colliculus. We recorded simultaneously from neurons in lateral intraparietal cortex (LIP) and the superior colliculus (SC) while monkeys made perceptual decisions, reported by eye-movements. Single-trial analyses revealed distinct dynamics: LIP tracked the accumulation of evidence on each decision, and SC generated one burst at the end of the decision, occasionally preceded by smaller bursts. We hypothesized that the bursts manifest a threshold mechanism applied to LIP activity to terminate the decision. Focal inactivation of SC produced behavioral effects diagnostic of an impaired threshold sensor, requiring a stronger LIP signal to terminate a decision. The results reveal the transformation from deliberation to commitment.

## Introduction

Decisions are elemental to almost all behaviors. Innate behaviors, such as escaping a predator, and our most complex behaviors, such as choosing a career path, involve similar processes—an evaluation of evidence for a set of options and a subsequent commitment to a proposition or plan of action. Much progress has been made in understanding how the brain evaluates or accumulates evidence during the formation of decisions. Less is known about how this process is terminated and how its outcome is transformed into a relevant plan or action.

For difficult decisions informed by noisy evidence, it is often useful to accumulate many samples of evidence until some stopping criterion is achieved. Such processes comprise a class of bounded random walks and drift diffusion models (Deco et al., 2013; Smith and Ratcliff, 2004; Gold and Shadlen, 2007). In monkeys trained to communicate their decisions with an eye movement, neurons in the lateral intraparietal area (LIP) represent the accumulation of evidence as the decision is being formed (Roitman and Shadlen, 2002; Huk and Shadlen, 2005). In addition, these neurons reach a stereotyped firing rate just before the decision is reported, which has led to the hypothesis that downstream areas—particularly those involved in generating the eye movement—apply a threshold on LIP activity and terminate the decision process when that threshold is exceeded (Roitman and Shadlen, 2002; Lo and Wang, 2006; Hanes and Schall, 1996).

A primary downstream target of LIP is the superior colliculus (SC), a conserved midbrain structure involved in orienting behaviors (White and Munoz, 2011). In primates, SC plays a prominent role in the generation of eye movements via its descending projections to brainstem oculomotor nuclei (Gandhi and Katnani, 2011). SC also sends ascending projections, via the thalamus, to the basal ganglia and cerebral cortex, including area LIP (Krauzlis et al., 2013). Lo and Wang (2006) proposed on theoretical grounds that SC is well-positioned to implement the decision threshold when decisions are communicated with an eye movement. However, experimental evidence for this proposal is lacking. Previous studies have shown that SC activity is qualitatively similar to LIP activity (Horwitz and Newsome, 2001b; Jun et al., 2021; Cho et al., 2021), supporting the idea that both SC and LIP represent the accumulation of evidence. Nevertheless, the hypothesis that SC applies a decision threshold on LIP activity has not been tested directly, primarily because it is challenging to record simultaneously from neurons in LIP and SC with the same spatial selectivity. The advent of a new generation of high-density multi-channel electrodes now renders such experiments possible.

Here, we provide evidence that SC implements the decision threshold. We recorded simultaneously from populations of neurons in LIP and SC that share the same spatial preference while monkeys performed a reaction-time, motion discrimination task. Single-trial dynamics in LIP approximate a stochastic drift-diffusion signal, consistent with the accumulation of noisy evidence. In contrast, single-trial dynamics in SC display bursts of activity, which terminate the decision. Simultaneous recordings suggest that these bursts reflect the implementation of a threshold applied to the drift-diffusion signal in LIP. We show that focal inactivation of SC impairs such implementation, leading to slower, biased decisions and prolonging the accumulation of evidence in LIP.

## Results

Two rhesus monkeys performed a reaction-time (free response) version of a commonly used motion discrimination task (Newsome et al., 1989; Roitman and Shadlen, 2002). The task requires the monkey to decide the net direction—left or right—of a small patch of dynamic random dots (Fig. 1A). After stimulus onset, the monkeys were free to report their decision with an eye movement to the leftward or rightward choice target. The direction and strength of the random dot motion (RDM) on each trial was selected pseudorandomly. We used a signed motion coherence to quantify the strength and direction, where positive values indicate rightward motion.

**Figure 1:**
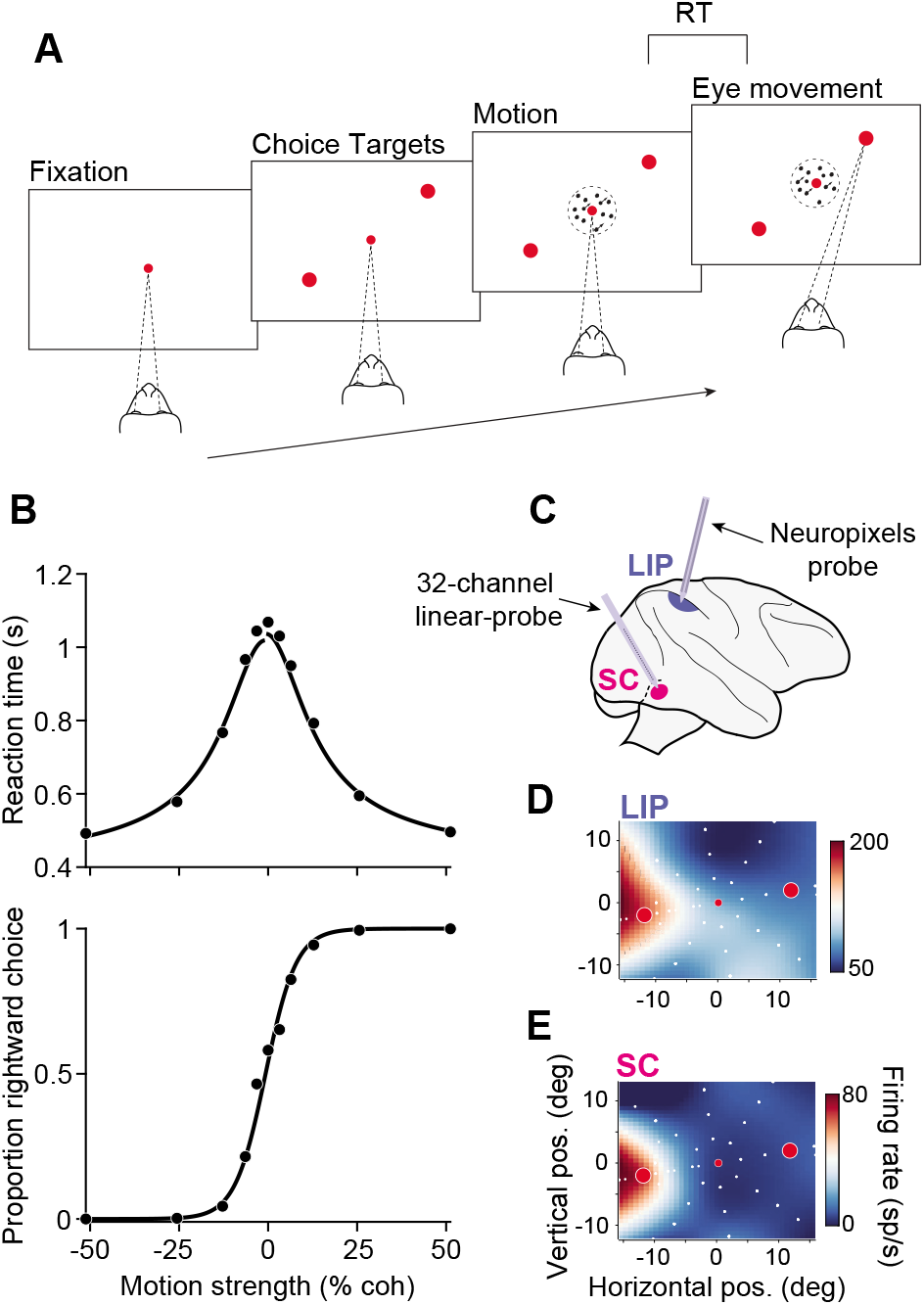
Behavioral task and experimental setup. **A,** Reaction-time (RT) motion discrimination task. Upon fixation and onset of the choice targets, a random-dot-motion stimulus appeared in the center of the display. Two monkeys discriminated the direction of motion and, when ready, indicated their choice with a saccade to one of two choice-targets. **B**, Behavioral data pooled across all sessions and both monkeys (19,749 trials). RT *(top)* and the proportion of rightward choices *(bottom)* are plotted as a function of motion strength. Positive (negative) motion strengths indicate rightward (leftward) motion. Solid curves depict fits of a bounded evidence accumulation model. **C**, Simultaneous recordings in SC and LIP. Populations of neurons were recorded in SC with a 16-32 channel V-probe and in LIP with a neuropixels probe. **D**, the response field (RF) of an example LIP neuron. The colormap depicts the mean firing rate, interpolated across target locations (white circles), during the delay epoch of a visually-guided saccade task. Red circles depict the location of the choice targets in the RDM task in this session. **E**, The RF of an example neuron in SC, recorded simultaneously as the example LIP neuron in **D**.

The monkeys’ choices and reaction times (RT) depended systematically on motion strength and direction (Fig. 1B). Both monkeys made no errors on the strongest motion (*C* = ±51.2%) and approached chance performance on the weakest stimuli (*C* ≈ 0%). Mean reaction times were less than a half second for the strongest motion strengths and more than a second for the weaker motion strengths. These observations are broadly consistent with predictions of bounded evidence accumulation models, in which the decision is terminated when a threshold level of accumulated evidence for leftward or rightward is exceeded. A fit of such a model to the behavioral data is shown by the black curves in Fig 1B.

The effect of 100 ms pulses of weak motion on choice and RT (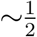 of trials) further supports that the animals decisions were based on accumulated evidence. The direction (left or right) and time of the pulse was chosen randomly, independent of the direction or strength of the stimulus (see methods). Figure S1 shows that the pulses affected choices and reaction times. Moreover, pulses shown early in the trial affected choices made nearly a second later. This persistent influence suggests that the monkeys’ decisions arise through the temporal integration of motion evidence and is inconsistent with non-integration strategies (Stine et al., 2020; Ditterich, 2006; Cisek et al., 2009). Together, the behavioral data suggest that the monkeys accumulated evidence over time and terminated their decisions when a threshold level of accumulated evidence was exceeded.

### Single-trial analyses reveal different dynamical processes in LIP and SC

We recorded from single neurons in LIP and SC while the monkeys performed the RDM task. We targeted the intermediate and deep layers of SC, where LIP afferents terminate (Paré and Wurtz, 1997; Lynch et al., 1985; Andersen et al., 1990), using 16-32 channel linear probes. Each session yielded 13-36 SC neurons, most of which had similar response field (RF) locations. During the RDM task, one choice-target was placed where overlap of RFs was maximal; the other choice-target was placed in the opposite hemifield (Fig. 1D). In LIP we used a prototype, neuropixels probe optimized for use in macaques (Neuropixels 1.0-NHP45), yielding 54-203 neurons per session. Of these neurons, we identified subsets of 9-34 that, by chance, had RFs that overlapped those of the simultaneously recorded SC neurons (Fig. 1D-E). We focused our analysis on these spatially-aligned neurons because previous work has shown that SC-projecting neurons in LIP share similar spatial selectivity to their target neurons in SC (Paré and Wurtz, 1997, 2001).

Consistent with previous results, we found that LIP and SC display similar trial-averaged activity during the RDM task (Fig. 2A,B). Activity in both areas is modulated by the motion strength and direction, and both predict whether the trial will end in a leftward or rightward choice. The most salient difference between LIP and SC activity is the large burst of activity in SC that begins approximately 150 ms before the saccade. The areas differ in their response to the brief motion pulses. Fig. 2C&D display the effect of pulses on activity in LIP and SC after controlling for other factors (Eq. 8). Consistent with temporal integration, pulses affect LIP activity for several hundred milliseconds (σ =.11 s, Gaussian fit). In contrast, the pulses affect SC activity more transiently (σ = .03 s; *p* < .001, likelihood ratio test), and this holds when the analysis is restricted to visuomovement prelude neurons identified by spatially-selective persistent activity (Fig. S2).

**Figure 2:**
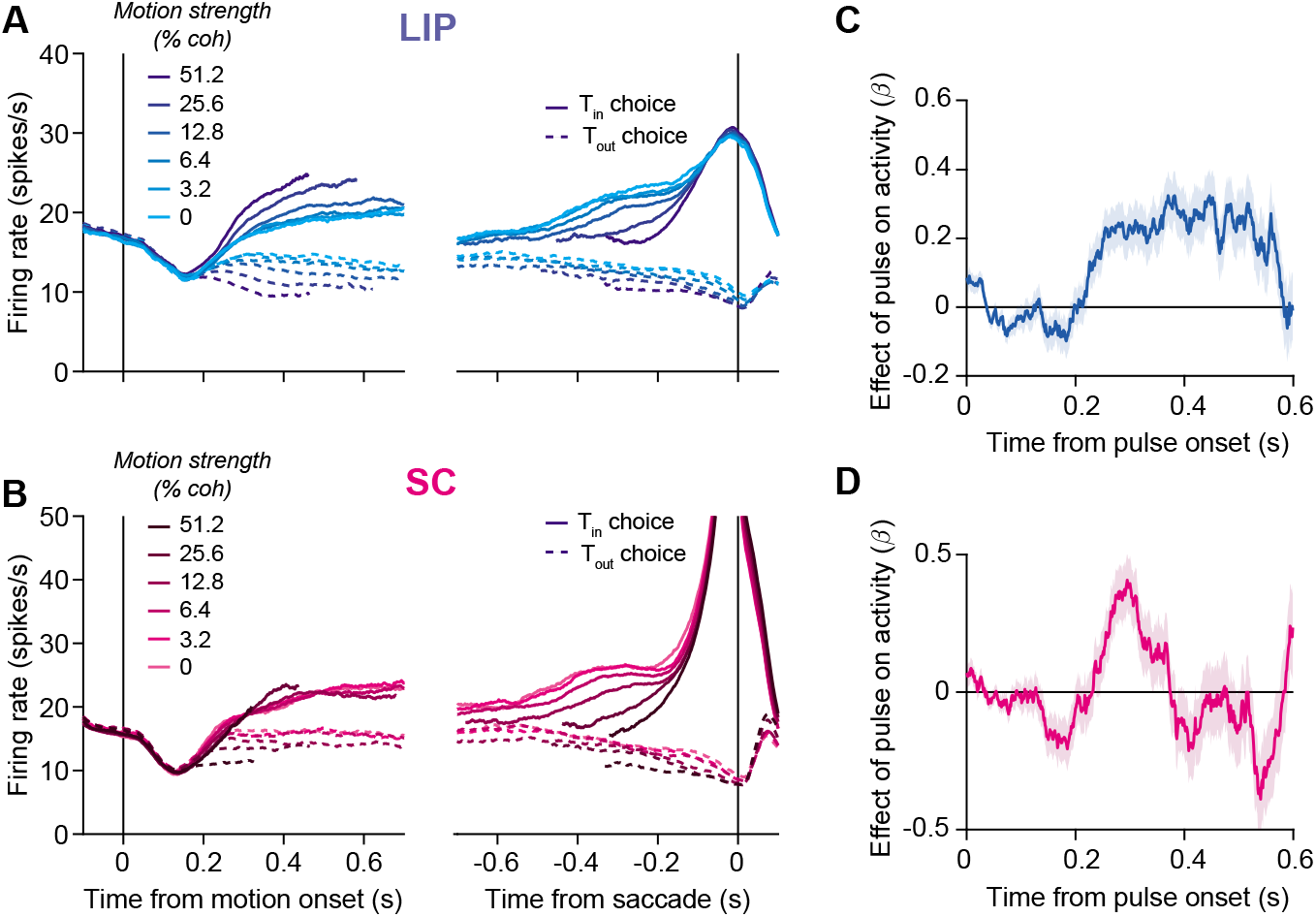
Trial-averaged responses of neurons in LIP and SC. **A,** Average firing rate of 164 neurons in LIP from two monkeys, aligned to motion onset *(left)* and saccadic onset *(right).* Responses are grouped by motion strength (shading) and direction (line style). Only correct trials are shown for non-zero coherences. **B,** Same as in **A**, but for 119 neurons in SC. **C**, Effect of motion pulses on LIP activity. Consistent with temporal integration, pulses had a persistent effect on LIP activity. Shaded region represents SE. **D**, Effect of motion pulses on SC activity. Pulses had a more transient effect on SC activity.

The contrasting effect of motion pulses on activity in LIP and SC is consistent with the hypothesis that the two areas perform different computations. To test this further, we exploited the fact that we recorded from many neurons in each area simultaneously. These large-scale recordings allowed us to observe single-trial dynamics in each area. Fig. 3 shows single-trial activity from example sessions in LIP and SC, aligned to motion onset and to the saccade. Each trace represents the average of the LIP or SC population on a single trial. The upper plots depict trials from 0% coherence trials, the middle plots depict trials from an intermediate motion strength (12.8% coherence), and the lower plots depict trials from the strongest motion strength (51.2% coherence).

**Figure 3:**
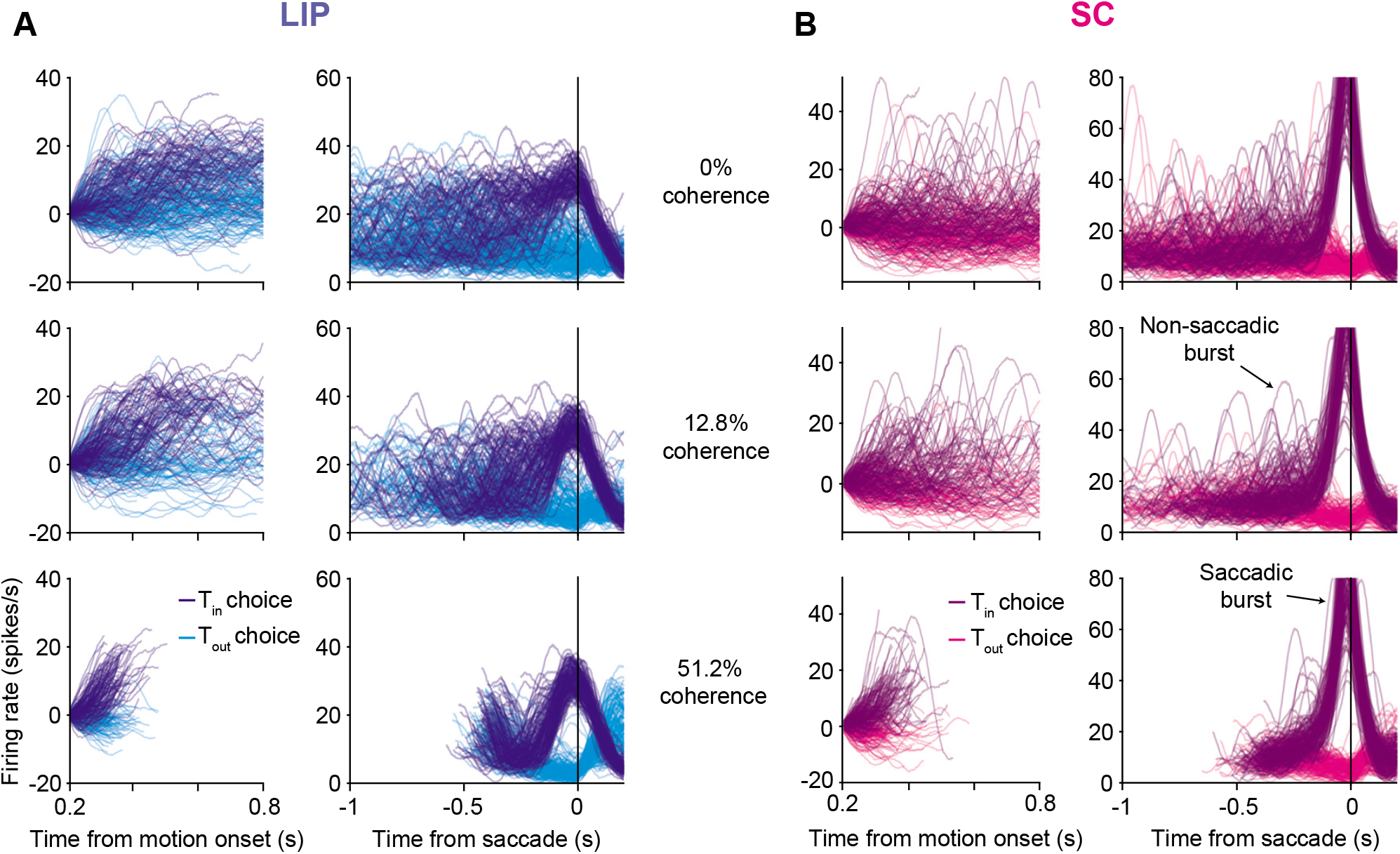
Single-trial dynamics in LIP and SC are different. **A,** Single trial activity in LIP from of an example session. Each trace depicts the average of 17 LIP neurons on a single trial, after smoothing spike trains with a Gaussian kernel (σ = 25 ms), for three different motion strengths (rows). The line color indicates the animal’s choice on that trial. Both correct and error trials are included. *Left column:* activity aligned to the onset of decision-related activity, ~200 ms from motion onset, until 100 ms before the saccade or 800 ms after motion onset (whichever occurs first). The rates are offset by an amount to force all traces to begin at zero. *Right column:* the same trials are shown aligned to saccade initiation, without baseline offset. Note the different ordinate scale. **B**, Same as in **A** but for 10 SC neurons. Two types of bursts were identified in SC: Saccadic bursts occurred at the end of the decision, just before the saccade; Non-saccadic bursts occurred as the decision was ongoing.

Single-trial responses in LIP (Fig. 3A) approximate drift-diffusion—the accumulation of noise plus signal. The ramp-like trajectories in the trial-averaged data (Fig. 2) reflect the accumulation of the signal (deterministic drift), whereas the accumulation of noise (diffusion) is suppressed by averaging. It is only evident in the single trial averages. In an accompanying paper, we show that such single trial firing rates establish the decision variable that determines the choice and RT on each decision (see Steinemann et al., 2022, for details).

Single-trial activity in SC is qualitatively different. In addition to the large saccadic burst, single trials elucidate additional bursts of activity as the decision is being formed. These bursts resemble the saccadic bursts, but they are not associated with a saccade and exhibit smaller amplitudes (Fig. S3). They are not apparent in the trial-averaged data because they are not aligned temporally to any trial-event. We developed an algorithm that classifies a high firing-rate-event as a burst if its derivative exceeds a positive threshold (see Methods). We found that non-saccadic bursts occurred in SC on 10.4% of trials. Crucially, they cannot be explained by the occurrence of small eye movements such as microsaccades (Fig. S3). Based on the distinct dynamics in the two areas, we hypothesized that SC applies a threshold on LIP activity, manifesting as a burst when exceeded. Typically, the burst terminates the decision, but it sometimes fails to do so.

### A threshold computation in the superior colliculus

Simultaneous recordings reveal a relationship between single-trial activity in LIP and bursting in SC. Fig. 4A shows the average LIP activity aligned to the onset of saccadic and non-saccadic bursts. Around the onset of the saccadic burst, LIP activity increases sharply and reaches a peak at the onset of the saccade. This “uptick” in activity immediately before a saccade is a known feature of LIP neurons; they are also evident in the saccade-aligned activity in Figs. 2 and 3. Note that these upticks have sometimes been termed a perisaccadic “burst” in previous work *(e.g.,* Paré and Wurtz, 2001). To avoid confusion, we will refer to “upticks” and “bursts”in LIP and SC, respectively.

**Figure 4:**
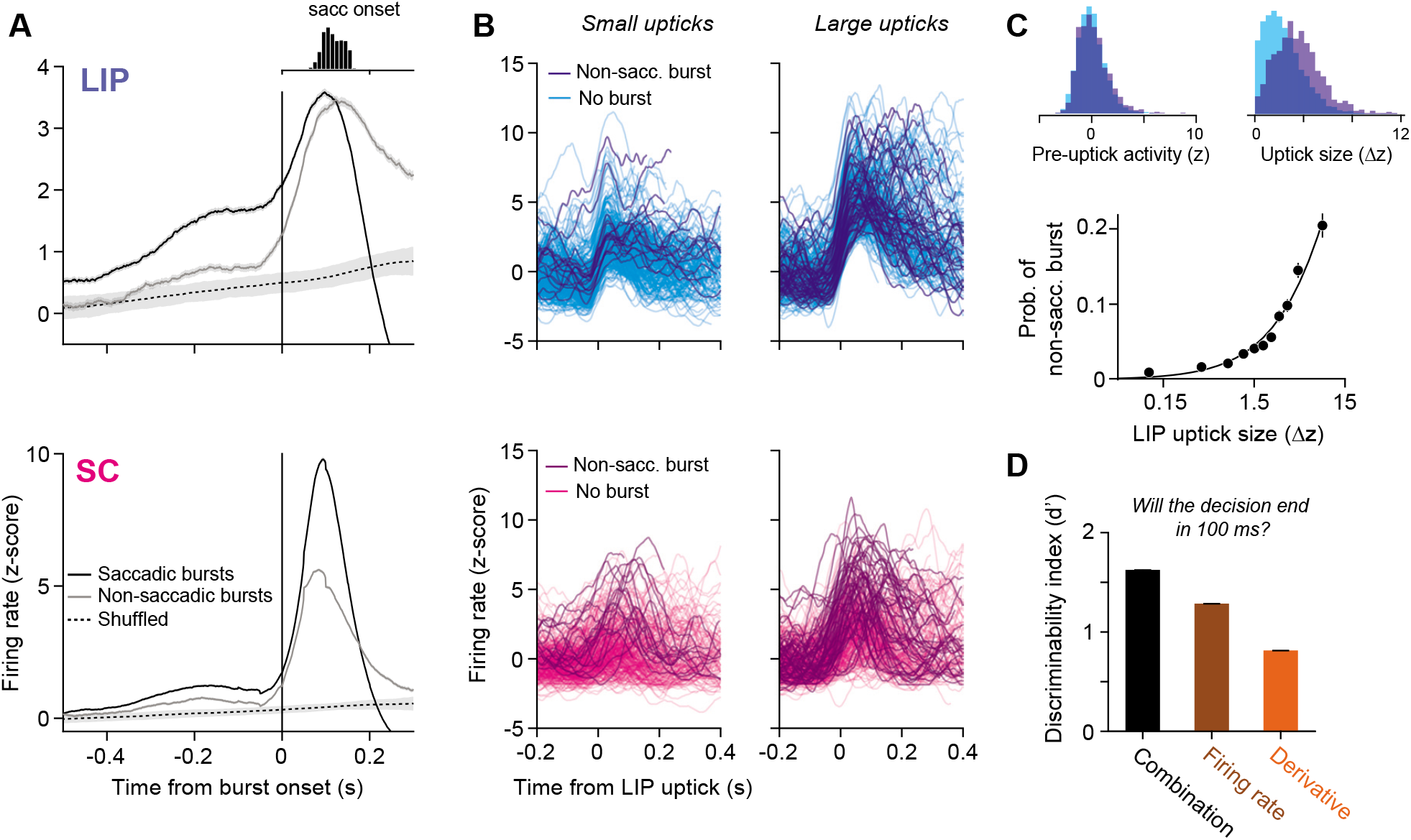
Upticks in LIP are associated with bursts in SC and decision termination. **A,** Average activity in LIP *(top)* and SC *(bottom)* aligned to the onset of bursts in SC or to random time points that preserve the temporal statistics of the non-saccadic bursts (shuffled, dashed line). Shaded region represents 95% CIs. Histogram *(inset)* shows the distribution of saccade initiation times relative to the onset of the saccadic burst (same time scale). **B**, Single trial firing rates aligned to small (*left*) and large *(right)* upticks in LIP activity. Darker traces represent trials in which a non-saccadic burst occurred in SC within 100 ms of the LIP uptick. **C**, Probability of a non-saccadic burst in SC increases as a function of the magnitude of the LIP uptick. *Top left* shows the distribution of baseline firing rates preceding an uptick, split by whether an uptick was associated with a non-saccadic burst in SC (indigo) or not (cyan); *top right* shows the same, but for the distribution of uptick magnitudes. **D**, Discriminability index (d’) for three quantities that use the firing rate (brown), its derivative (orange), or a combination of the two (black) to predict whether the decision will terminate in 50-150 ms. Error bars (barely visible) represent standard error.

Similar upticks in LIP activity occur at the onset of non-saccadic bursts (Fig. 4A). However, instead of dropping precipitously within 200 ms, mean activity peaks and then remains elevated. In other words, the uptick in LIP activity that occurs before a saccade also occurs in association with non-saccadic bursts. Additional analyses suggest that non-saccadic bursts in SC are predicted by the size of the uptick in LIP. On each trial, we identified upticks in LIP activity and quantified their magnitude. Fig. 4B shows example upticks and the corresponding SC activity. The darker traces depict trials in which a non-saccadic burst occurred in SC within 100 ms of the LIP uptick. Non-saccadic bursts more commonly occur when upticks in LIP are large (Fig. 4C). Indeed, non-saccadic bursts occurred over 20% of the time during the largest LIP upticks, whereas they rarely occurred within a randomly-chosen, 100 ms window (2.1%). The observation that LIP upticks occur during both types of SC bursts suggests that the termination mechanism might be more complicated than a simple threshold on the LIP firing rate. Note that these analyses are correlational and therefore do not provide evidence that upticks in LIP cause bursts in SC, but we provide evidence below that the SC bursts do not cause upticks in LIP.

The distinction between upticks that are associated with a saccade and those that are not is that the former occur on top of an elevated firing rate (Fig. 4A). The result raises the possibility that upticks in LIP activity trigger bursts in SC and that saccadic bursts are triggered specifically when the uptick stems from a high firing rate. Indeed, such events accurately predict when the decision will terminate. We attempted to predict, at each moment (t) in each trial, whether the decision would terminate in a T_in_ choice 50-150 ms later. We applied a criterion to one of three quantities derived from the LIP activity: (1) the firing rate, *r*(*t*), (2) its derivative, 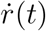, or (3) a weighted sum of the derivative and the firing rate from 50 ms earlier, 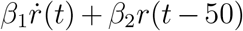. The third quantity captures the proposal that the combination of an uptick stemming from a high firing leads to decision termination. Using signal detection theory, we compared the capacity of the three quantities to discriminate, at each moment t, whether the decision will terminate. “Hits” are defined as time points in which the quantity of interest exceeds a criterion and a T_in_ saccade is initiated 50-150 ms later. “False alarms” are defined as time points in which the criterion is exceeded and the decision does not terminate in a T_in_ saccade (see Methods). Fig. 4D shows the discriminability index (d’) associated with each quantity (see Fig. S4 for ROC curves). The discriminability of the third quantity is highest 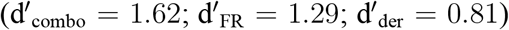, consistent with our proposal, and a nested model comparison suggests that the added degree of freedom is justified (*p* < 0.01, likelihood ratio test).

From these observations, we conclude that upticks in LIP, bursts in SC, and the subsequent termination of the decision are closely related to one another. We hypothesized that SC implements the decision threshold associated with a T_in_ choice, and it does so by sensing the combination of two events in LIP: an uptick and a high firing rate. Exceeding this threshold triggers a saccadic burst and terminates the accumulation process in LIP.

### SC plays a causal role in terminating decisions

If the hypothesis is correct, then inactivation of SC should impair the terminating threshold. We unilaterally inactivated SC with small volumes of muscimol, a GABA agonist, while recording simultaneously from neurons in LIP with a neuropixels probe (Fig. 5A). In each session, we first characterized the effect of inactivation on visually-instructed delayed saccades. Consistent with previous studies (Quaia et al., 1998; McPeek and Keller, 2004; Lovejoy and Krauzlis, 2010; Bollimunta et al., 2018a), focal SC inactivation increased saccadic latencies for saccades confined to a particular location in space, which we term the inac-tivation field (Fig. 5B). During the RDM task, one choice target was placed in the center of the inactivation field (T_in_); the other was placed in the opposite hemifield (T_out_).

**Figure 5:**
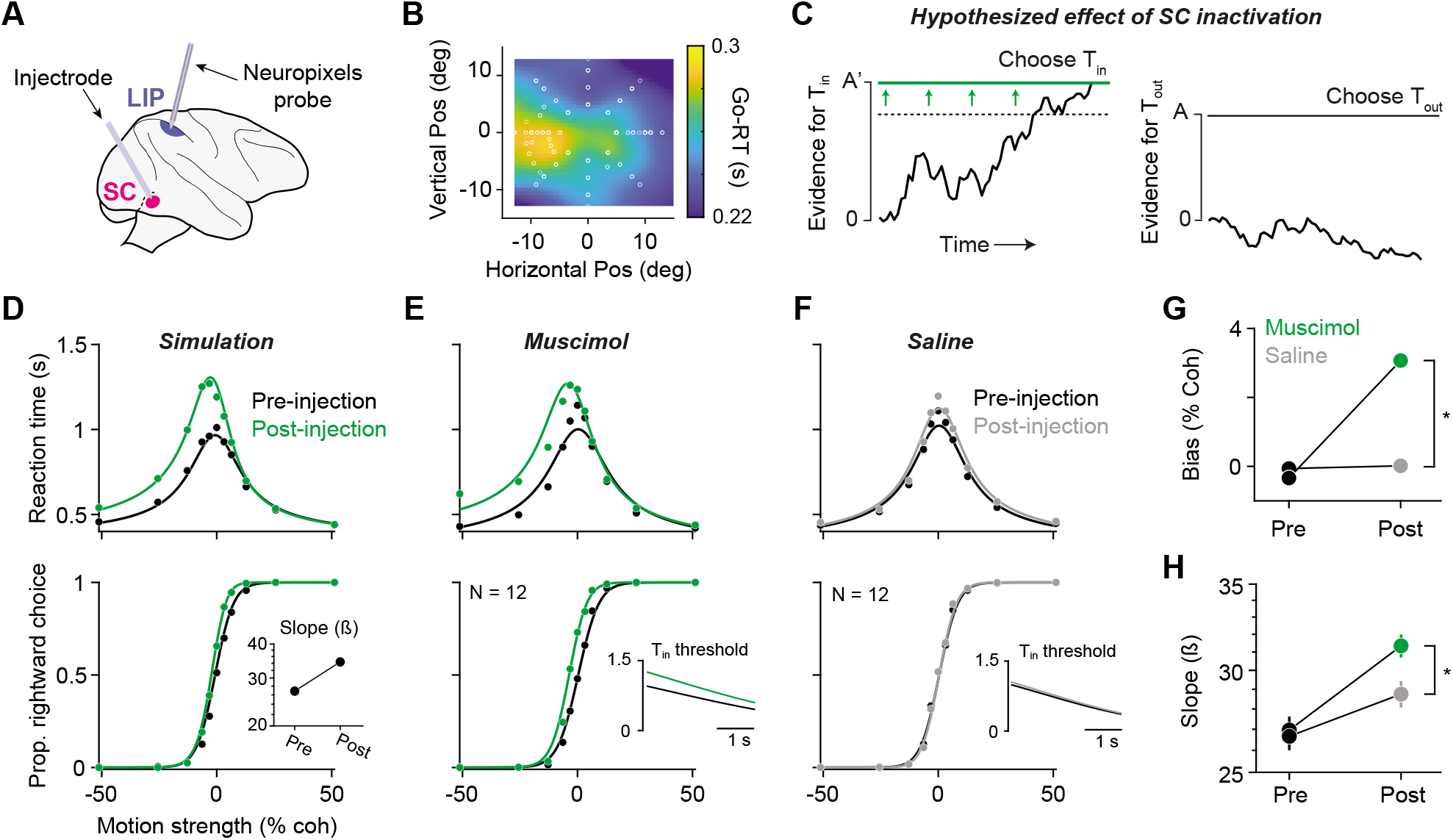
Focal inactivation of SC impairs decision termination. **A,** Experimental setup. Muscimol was injected unilaterally in the intermediate and deep layers of SC. On half of the experiments, Neuropixels recordings were obtained from ipsilateral area LIP. **B**, Saccadic latencies measured in an instructed delayed saccade task were slowed by 50 ms relative to pre-injection in a region of the contralateral visual field (heatmap). The left-choice target was placed in the center of this region. **C**, An impairment in the mechanism for detecting the threshold might result in a requirement for stronger signals to achieve termination for T_in_ choices. This is equivalent to application of higher decision threshold for T_in_ choices (left panel), while leaving the threshold for T_out_ choices unchanged (right panel). **D**, Simulated choice *(bottom)* and RT (*top*) data generated by the model in **C**, before (black data points) and after (green data points) a 70% increase in the Tin decision threshold. Curves show the fit of a bounded evidence accumulation model. *Inset* depicts the predicted effect of SC inactivation on the slope of the choice function. **E**, Choice-RT data before (black) and after (green) unilateral SC inactivation. *Inset* depicts the model-derived T_in_ decision threshold before and after SC inactivation. SC inactivation increased the T_in_ decision threshold by 31.9%. **F**, Same as **E** but for saline/sham injection experiments. **G**, Effects of muscimol (green) and saline/sham (grey) injection on choice bias. Positive values indicate a bias toward ipsilateral (rightward) choices. Asterisk denotes a statistically significant difference *(p* < .01, likelihood ratio test). **H**, Effects of muscimol and saline/sham injection on the slope of the choice function.

To develop an intuition for how a disrupted termination mechanism might affect decisions, it is useful to depict the decision process as a race between two accumulators that accumulate evidence for a leftward and rightward choice, respectively (Fig. 5C). In this architecture, the decision (leftward or rightward) is determined by the accumulator that first exceeds its corresponding threshold. LIP activity in the right hemisphere, say, corresponds to the leftward accumulator. According to our hypothesis, the decision threshold resulting in a leftward choice is implemented by the SC in the right hemisphere. Inactivation of the right SC should therefore interfere with the ability to commit to a T_in_ (left) choice. If the mechanism of termination is impaired, such commitment might require stronger signals from LIP, such as larger upticks, higher firing rates, or both. Conceptually, any of these changes would equate to an increase in the level of accumulated evidence required to trigger a T_in_ choice—that is, an increase in the T_in_ decision threshold (Fig. 5C). Such an increase would give rise to three observations in the behavior. First, because there is an asymmetry in the amount of evidence required for each choice, the monkeys should be biased away from T_in_ choices. Second, reaction times for T_in_ choices should increase, because it takes longer to exceed the increased T_in_ threshold. Third, the slope of the choice function *(i.e.* sensitivity) should increase following SC inactivation. The intuition for this last prediction is that the increased decision threshold causes decisions to be based on more accumulated evidence, which leads to better performance. (Fig. 5D) depicts these effects in simulated data, generated by a bounded evidence accumulation model in which the Tin decision threshold was increased by 70%.

We observed all three predicted effects in the monkeys’ behavior following unilateral SC inactivation. Fig. 5E displays choice and reaction time data before and after muscimol injection, combined across all sessions. SC inactivation caused a bias toward T_out_ choices (Fig. 5G), an increase in reaction times on T_in_ choices, and an increase in the slope of the choice function (Fig. 5H). All three effects were consistent across sessions and monkeys (Fig. S5). Additionally, the increase in contralateral reaction times is not fully accounted for by the increase in saccadic latency observed in the delayed saccade task (mean ΔRT = 0.18 s; mean ΔSL = 0.05 s). There was a general trend in which the slope of the choice function increased as the session went on, and indeed, this increase can be seen in the saline sessions. However, the increase in slope following muscimol injection is significantly larger than that after saline injection (p = 0.006, log-likelihood ratio test). While there are many potential mechanisms that might produce a choice bias and an increase in reaction times, only an increase in the decision threshold parsimoniously explains all three effects.

The conclusion is also supported by a formal model comparison. We fit choice-RT data using a drift-diffusion model with collapsing and asymmetric decision thresholds. We allowed the model to fit three extra parameters to capture the effect of SC inactivation on behavior: (1) a change in the T_in_ decision threshold, (2) a change in the T_out_ decision threshold, and (3) a shift in the drift-rate offset (see Methods for details). Fits of this model are shown by the solid curves in Fig. 5E, F. The insets display the model-derived T_in_ decision threshold before and after muscimol injection (E) or saline injection (F). These fits suggest that muscimol injection increased the T_in_ decision threshold by 31.9%, whereas it increased 6.6% following injection of saline. We also found that the effect of SC inactivation could not be fully explained by an increase in the Tin decision threshold. A significant shift in the drift-rate offset was also ascribed by the model, which is consistent with a previous study of SC inactivation (Jun et al., 2021). Together, the effects of focal SC inactivation on behavior provide evidence that SC plays a causal role in terminating the decision process when a threshold level of accumulated evidence is exceeded.

### The altered decision threshold is apparent in LIP activity

If this interpretation is correct, we might expect SC inactivation to distort the events in LIP that predict decision termination. For example, decision termination might require larger upticks, higher firing rates, or both. In half of the muscimol sessions, we recorded activity in LIP with a neuropixels probe and identified LIP neurons with response fields that overlapped the choice target in the inactivation field. As shown in Fig. 6A, inactivation of SC led to an increase in LIP firing rate in the epoch preceding decision termination (FR_pre_ = 20.6, FR_post_ = 26.4), and the increase in activity over time is more gradual. The observation is present in most neurons Fig. 6B, and it is not explained by a more general increase in firing rates at all time points or an increase in gain. Indeed, SC inactivation reduced the magnitude of visual responses to the choice targets and caused a subtle decrease in build-up rates early in the trial (Fig. S6). We did not observe an increase in LIP activity following saline injection (Fig. S6).

**Figure 6:**
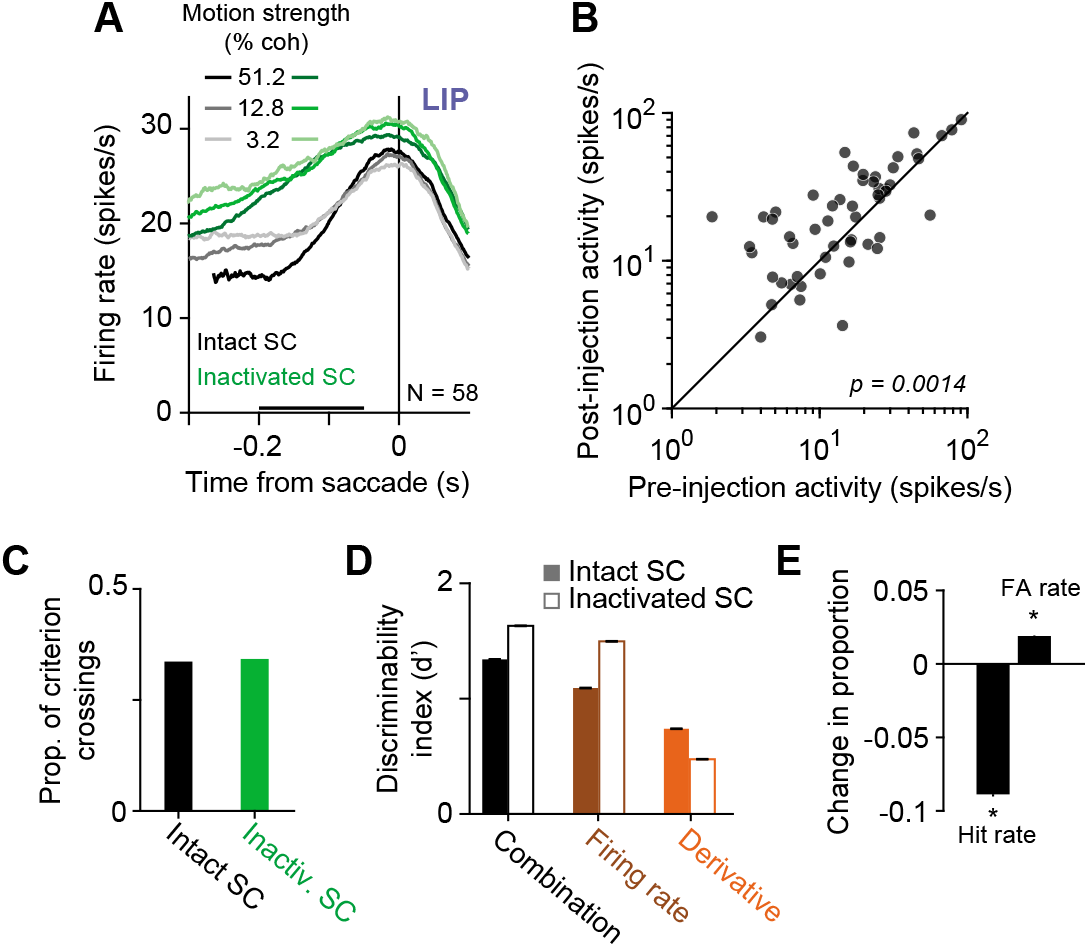
Altered read-out of LIP activity following SC inactivation. **A,** Effect of SC inactivation on LIP population activity at the end of the decision. Mean firing rates of 58 LIP neurons before (grey curves) and after (green curves) SC inactivation are aligned to the onset of the saccade. Black bar denotes the epoch used for the analysis in *B.* **B,** Comparison of mean firing rates of individual neurons before and after SC inactivation. Activity was significantly greater after SC inactivation *(p* = 0.001, paired t-test). **C**, The proportion of time points in which the firing rate derivative exceeded a criterion before (black) and after (green) SC inactivation. The analysis suggests that SC inactivation did not reduce the frequency or magnitude of upticks in LIP. **D**, The same analysis as in Fig. 4D applied to data before and after SC inactivation. Note the decrease in d’ associated with the derivative signal and the increase associated with the other two quantities. **E**, Change in hit rate and false alarm rate associated with the derivative after SC inactivation, assuming no change in criterion. Asterisk denotes p < 0.05, chi-squared test.

Finally, we found that SC inactivation disrupted the relationship between upticks in LIP and decision termination. After SC inactivation, the uptick associated with the saccade was less apparent in the trial-averaged traces (Fig. 6A). The observation could arise if LIP upticks do not occur after SC inactivation. However, this is not the case. We applied a criterion to the derivative of the firing rate and computed the proportion of criterion crossings before and after SC inactivation. Fig. 6C shows that the derivative achieves this criterion with an almost identical frequency before and after SC inactivation (*P_pre_* = 0.336, *P_post_* = 0.342). Instead, the upticks in LIP were less predictive of decision termination. We repeated the analysis in Fig. 4D in which we use LIP activity to predict when the decision will terminate. The discriminability of the derivative signal, which captures whether upticks are predictive of decision termination, decreased after SC inactivation (Fig. 6D). In contrast, the discriminability of the other two quantities increased. The observation suggests a change in the downstream mechanism that senses upticks in LIP activity. Consistent with this, the derivative signal was associated with a higher false alarm rate and a lower hit rate after SC inactivation, assuming a constant criterion (Fig. 6E). In other words, upticks still occurred, but their correlation with decision termination was blunted.

## Discussion

We have identified a neural mechanism in the primate superior colliculus for terminating a decision based on the state of accrued evidence. It has been hypothesized that a threshold is applied to firing rates in association cortex to terminate decisions (Roitman and Shadlen, 2002; Lo and Wang, 2006; Hanks et al., 2014; Ding and Gold, 2012; Hanes and Schall, 1996; Heitz and Schall, 2012). Until now, the mechanism for how this is achieved was unknown. We show that neurons in LIP represent the accumulation of noisy evidence on single decisions—the latent drift-diffusion process posited by evidence accumulation models (see also Steinemann et al., 2022). SC, which is reciprocally connected with LIP, performs a distinct computation. It implements a threshold, manifesting as a burst, that generates a saccade and terminates the decision process. We show that inactivation of SC impairs this threshold mechanism, leading to longer decisions and prolonged accumulation in LIP. These insights were made possible by a new generation of high-density multi-channel electrodes—45 mm Neuropixels probes—which enable simultaneous recordings from many neurons with similar spatial selectivity to reveal firing rates of functionally similar neurons in the SC and LIP at the same time on a single decision.

Previous studies support the idea that SC and LIP represent similar decision-related signals (Horwitz and Newsome, 1999, 2001b,a; Ratcliff et al., 2003, 2007; Cho et al., 2021; Basso et al., 2021; Jun et al., 2021). Based on trial averaged activity, both SC and LIP appear to represent the accumulation of evidence (Fig. 2). The averages in both areas exhibit evidence-dependent buildup (or decrease), sometimes referred to as ramping. We show that in LIP, these averages belie drift-diffusion signals on single decisions (Fig. 3A; see Steinemann et al. 2022). In stark contrast to LIP, the trial-averaged spike rates in SC belie non-saccadic bursts. These bursts do not account fully for the trial averages in SC, but they are the most salient feature. It is possible that non-saccadic bursts contributed to the trial-averaged activity in previous studies of SC. For example, Cho et al. (2021) reported step-like activity in a detection task and Horwitz and Newsome (2001b) found evidence for large transitions in single-trial spike trains. Our results do not preclude the possibility that small subsets of SC neurons, such as those with direction selectivity (Horwitz and Newsome, 2001a), display single-trial dynamics that match those of LIP. Given the scarcity of such neurons, resolving this question would require many more simultaneously recorded neurons. Nevertheless, our results show that neurons in SC and LIP perform different computations despite their many similarities.

Through simultaneous recordings, we provide evidence that SC applies a threshold to LIP activity using a combination of the firing rate and its derivative. Although we do not know whether the neurons recorded in LIP and SC are connected directly, they are likely representative of—and correlated with—those that are, owing to their overlapping spatial preferences (Paré and Wurtz, 1997). The observation that upticks are associated with both non-saccadic and saccadic bursts may provide insight into the bursting mechanism within SC (Isa and Hall, 2009). The preservation of upticks when SC is inactivated suggests that LIP does not depend on the portion of inactivated SC to generate the upticks. The upticks themselves are part of the decision variable in LIP—the rising phases of drift diffusion signals (Steinemann et al., 2022).

The threshold mechanism we propose almost certainly involves cooperation among other visuomotor association areas, such as FEF and the basal ganglia. In particular, we speculate that the role of the substantia nigra pars reticulata (SNr), an output nucleus of the basal ganglia, is critical. SNr provides tonic inhibition to SC, which decreases before an eye movement (Hikosaka and Wurtz, 1983a,b). We envision a circuit mechanism with two properties: (1) a mechanism in SC that triggers a burst when an uptick is sensed, and (2) tonic inhibition of SC, via SNr, that is negatively correlated with the level of activity in LIP. When activity in LIP is weak, inhibition of SC is strong; thus, upticks sometimes trigger a burst in SC, but it is quickly suppressed and the decision continues. When activity in LIP is strong, inhibition of SC is weak, unleashing SC to generate a large saccadic burst if it receives an uptick from cortex. This circuit mechanism is similar to the model proposed by Lo and Wang (2006). A natural question is whether slight modifications to their model could explain our results or whether a different type of threshold mechanism would be required *(e.g.* Evans et al., 2018).

We confirmed that SC is causally involved in terminating decisions through the use of focal inactivation. Inactivation caused a bias against contralateral (Tin) choices, coupled with an increase in RT on those choices, and an increase in the slope of the choice function. The pattern is diagnostic of an increase in the Tin decision threshold. More accumulated evidence is required to commit to a Tin choice, because the termination mechanism is impaired. These effects of SC inactivation on decisions are distinct from those observed in studies that inactivated LIP. Unilateral inactivation of LIP led to an ipsilateral choice bias without the hallmarks of a change in termination threshold (Jeurissen et al., 2021; Zhou and Freedman, 2019) or no effect (Katz et al., 2016). In a recent study, Jun et al. (2021) found that unilateral inactivation of SC induced an ipsilateral choice bias. We uncover a potential neural correlate of this bias effect, manifesting as a slight decrease in LIP build-up rates. In contrast to our results, however, Jun et al. (2021) did not observe the hallmarks of an increased decision threshold. We speculate that their subjects may not have been terminating decisions at a threshold level of accumulated evidence (see their supplemental table 4), which would explain the discrepancy in our results.

SC and LIP are reciprocally connected. It is therefore natural to wonder if SC inactivation affects behavior through an effect on LIP. However, neural recordings from LIP during SC inactivation indicate that the effects are downstream of LIP The drift-diffusion signal in LIP was qualitatively unaffected—just prolonged, leading to higher firing rates at the end of the decision. We interpret this as a sign of an impaired termination mechanism, which requires a stronger signal. The need for a stronger LIP signal would hold whether the termination is computed by SC neurons outside the inactivation field (Lee et al., 1988) or by a mechanism that bypasses the SC altogether, for example, by exploiting the direct projection from FEF to oculomotor nuclei in the pons (Büttner-Ennever and Horn, 1997; Moschovakis and Highstein, 1994; Segraves, 1992). The higher firing rates at termination are not trivially explained by the fact that decisions are prolonged. Previous studies in which animals were encouraged to slow down their decisions found either the opposite effect (Heitz and Schall, 2012) or no change in activity at the end of the decision (Hanks et al., 2014).

The results are likely to extend to more common settings. In our task, the oculomotor system is used to convey a decision and extract a reward. In more natural settings, it is typically used to interrogate objects and locations in the world. Such interrogation is accomplished either overtly, via an eye movement, or through covert spatial attention. The role of SC in the former has long been appreciated, but a more recent body of work has revealed its critical role in the latter (Kustov and Robinson, 1996; Ignashchenkova et al., 2004; Bogadhi et al., 2021; Herman et al., 2020, 2018; Bollimunta et al., 2018b; Müller et al., 2005; Zénon and Krauzlis, 2012; Lovejoy and Krauzlis, 2010). We view our results and SC’s role in spatial attention as two sides of the same coin. Indeed, the allocation of attention is naturally framed as a decision process informed by sensory evidence. In LIP, the accumulated sensory evidence may take the form of a priority map of objects worth inspecting (Bisley and Goldberg, 2010), and the inspection is implemented, either covertly or overtly, when a burst occurs in SC.

Finally, distinct roles played by LIP and SC bear on a fundamental point about neural computation. LIP and SC are just two nodes in a network of brain regions that play a role in perceptual decisions reported by an eye movement. Based on average firing rates, it is tempting to conclude that the computations for forming and terminating decisions are distributed across these nodes, with no single area playing a specialized role. The present result supports a more modular organization for forming and terminating a decision. There is a certain appeal to such modularity. It allows for the possibility that some signals might affect the representation of evidence without influencing the decision threshold, or it might enable different effector systems to establish different thresholds. For example, it might be sensible to acquire more evidence to reach for something than to look at it. From an evolutionary perspective, the specific cortical-midbrain organization supports the idea that cognitive, deliberative decisions have simply co-opted primitive circuits that underlie innate perceptual decisions, such as orienting, freezing, and escape (Yilmaz and Meister, 2013), and their study may provide the answers to more fundamental questions about how the mechanism works at a biophysical level (Evans et al., 2018; Lo and Wang, 2006). Most work has focused on how the brain forms decisions, but an understanding of how the brain commits to a decision is just as critical. Indeed, deliberation is useless or—as in the case of Buridan’s donkey (Blackburn, 2008)— debilitating if it cannot be terminated.

## Methods

The data in this study were obtained from two adult male rhesus monkeys *(Macaca mulatta,* 8-11 kg; Monkey M and Monkey J). All training, surgery, and experimental procedures complied with guidelines from the National Institutes of Health and were approved by the Institutional Animal Care and Use Committee at Columbia University. A head post and two recording chambers were implanted using aseptic surgical procedures and general anesthesia. Placement of the LIP chamber was guided by structural MRI. The SC chamber was placed on the mid-line and angled 38°posterior of vertical. The experiment was controlled by the Rex system (Hays et al., 1982) on a QNX operating system. All visual stimuli were displayed on a CRT monitor (75 Hz refresh rate, 57 cm viewing distance) controlled by a Macintosh computer running Psychtoolbox (Brainard, 1997). Eye position was monitored with an infrared video tracking system with a 1 KHz sampling rate (Eyelink 1000; SR Research, Ottawa, Canada).

### Behavioral tasks

The monkeys performed two tasks in each session. In the *instructed-delay* saccade task (Hikosaka and Wurtz, 1983a), a visual target appeared in the periphery as the monkey fixated a central fixation point (FP). The monkey was required to maintain fixation until the FP disappeared, thereby allowing the monkeys to execute a saccade to the target. In a memory-guided variant of the task (Gnadt and Andersen, 1988), the target was flashed for 200 ms and the monkey was required to execute a saccade to its remembered location when the FP was extinguished. Like many intervals in our tasks, the delay period was determined by a random draw from a truncated exponential distribution:

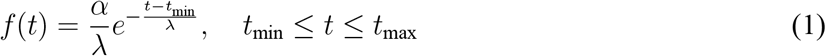

where *t*_min_ = 0.5 s and *t*_max_ = 1.5 s define the range, λ = 0.7 s is the time constant, and *α* is chosen to ensure the total probability is unity. Below, we report the range and the exponential parameter λ. Because of truncation, the expectation 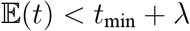.

The reaction-time (RT) motion discrimination task is similar to previous studies (*e.g.* Roitman and Shadlen, 2002). The monkey initiated a trial by foveating the FP, after which two choice-targets appeared on the screen (one in each hemifield). After a random delay (truncated exponential: 0.25-0.7 s; λ = 0.4 s; Eq. 1), a random-dot-motion (RDM) stimulus was displayed centrally, subtending 5 degrees of visual angle. Details of the RDM stimulus have been described previously (*e.g.* Britten et al., 1996; Roitman and Shadlen, 2002) and its spatial and temporal statistics were identical to those used in (Stine et al., 2020). The monkey was required to judge the direction (leftward or rightward) of motion embedded in the stimulus and report its decision by making a saccade to the leftward or rightward choice target. The strength of the motion *(i.e.* motion coherence) was chosen pseudorandomly on each trial and could take one of the following values: 0%, 3.2%, 6.4%, 12.8%, 25.6%, and 51.2%. After the onset of the RDM stimulus, the monkey was free to report its decision and RT was defined as the interval between the onset of the RDM stimulus and the initiation of the saccade. Correct choices were rewarded with a juice reward and were followed by an intertrial interval of 0.75 s. Incorrect choices were not rewarded and were followed by an additional time-out of 0-3 s. We included two incentives to discourage rushed responses. The first was that the time-out after an error trial was a function of RT, such that faster incorrect choices were followed by longer time-outs. The second was enforcement of a minimum time of 800 ms between motion onset and the reward. For example, a RT of 500 ms would lead to a delay of 300 ms before reward delivery, whereas RTs ≥ 800 ms would incur no delay.

On approximately half of the trials, a 100 ms pulse of motion was added to the RDM stimulus. The pulse took the form of an increment or decrement of motion coherence (similar to Kiani et al., 2008). For example, if the motion stimulus had a signed coherence of +12.8%, a leftward motion pulse of strength 4.0% would change this coherence to +8.8% for 100 ms. The strength of the pulse was calibrated to induce a weak but reliable effect on choices and RTs. The pulse strength was 4.0% coherence for Monkey M and 3.2% for Monkey J. The sign of the pulse was chosen randomly and was independent of the motion direction. The onset of the pulse was drawn randomly from a truncated exponential distribution (0.1–0.8 s, λ =0.4, Eq. 1).

### Simultaneous recordings

We recorded from well-isolated single neurons in LIP and SC using multi-channel electrodes. We targeted the ventral subdivision of LIP (Lewis and Van Essen, 2000), which is defined anatomically by its projections to the frontal eye field (FEF) and the intermediate and deep layers of SC (Andersen et al., 1990). We targeted these layers in SC. In LIP, most of the data were recorded with neuropixels probes optimized for use in macaques (Neuropixels 1.0-NHP45). These probes are 45 mm in length and contain 4,416 recording sites, 384 of which are selectable to record from at one time. We conducted five recording sessions with the Neuropixels probe in Monkey M and three sessions in Monkey J, yielding 54-203 LIP neurons per session. These 8 sessions are also included in a companion paper focusing on the drift-diffusion signal in LIP (Steinemann et al., 2022). Neural data from these probes were recorded using SpikeGLX software and were synced post-hoc to behavioral data, task events, and SC activity recorded with an Omniplex system (Plexon). In Monkey M, we conducted an additional six sessions in which we used a 16-channel V-probe (Plexon) in LIP. All but one of these were not included in the data due to a lack of LIP neurons with an RF that overlapped those of the SC neurons. We recorded simultaneously in the ipsilateral SC with 16-, 24-, and 32-channel V-probes (Plexon; 50-100 μm electrode spacing), yielding 13-36 neurons per session.

Each session proceeded as follows: We first lowered the SC probe and measured the approximate response fields (RFs) of the SC neurons using the delayed saccade task. These RFs typically overlapped with one another due to our trajectory being approximately normal to the retinotopic map in SC. We only proceeded if the center of the RFs had an eccentricity greater than 7°of visual angle, which ensured that the RFs would not overlap the RDM stimulus. The intermediate and deep layers were identified by the progression from purely visual responses to motor and prelude responses as the probe advanced deeper. Once the location of the SC RFs was confirmed, we lowered the neuropixels probe into LIP and allowed for 15-30 minutes of settling time in order to maximize recording stability. The monkey then performed 100-500 trials of the delayed saccade task, with a variety of target locations, in order to precisely measure the RFs of the SC and LIP neurons. Finally, the monkey performed the RDM task until satiated—typically 1,500-3,000 trials— with a few dozen trials of the delayed saccade task included intermittently in order to measure stability of the RFs. In the RDM task, the location of the contralateral choice target was placed in the center of the SC neurons’ RFs. The other choice target was placed in the opposite hemifield, such that the two choice targets and the fixation point were collinear. The LIP neurons with RFs that overlapped the contralateral choice target were identified post-hoc by analyzing activity in the delayed saccade task (see below).

### SC inactivation experiments

We unilaterally inactivated SC with small volumes of muscimol (0.25-0.4 μL, 5 μg/μL), a GABAA receptor agonist. To target the intermediate and deep layers of SC, we recorded neural activity at the tip of a custom-built injectrode (30ga or 33ga cannula, similar to Chen et al., 2001) with a glass-coated tungsten microelectrode (Thomas Recording). The injection site was chosen to be at a depth that yielded strong motor activity, typically 2-2.5 mm below the surface of SC. The drug was injected with a syringe pump (Harvard Apparatus) connected to a 2 μL or 5 μL Hamilton syringe. Before lowering the injectrode, the monkey performed approximately 100 trials of the delayed saccade task and 800-1,000 trials of the RDM task to establish the pre-injection, baseline behavior. Following muscimol injection, we waited 15-30 minutes before testing for a deficit in saccade metrics. To assess the inactivation field, we measured saccadic latency and peak velocity for an array of target locations in the instructed-delay saccade task. The presence of an effect on saccade metrics was used as a positive control, which informed whether the injection was successful; the experiment was terminated if there was not a clear, spatially confined increase in saccadic latency or decrease in peak velocity following muscimol injection. Following identification of the inactivation field, the monkey performed the RDM task until satiated. In the RDM task, the contralateral choice target was placed in the center of the inactivation field. In two sessions in Monkey M, the inactivation field extended foveally, which caused a large increase in fixation breaks (>10% of trials). These sessions were removed from the dataset because of the worry that an increase in fixation breaks can affect decision strategy (see Chapter 3 of Kira, 2014). Saline injections followed the same protocol as that described above. In sham experiments, the injectrode was lowered into SC or just above it, but no solution was injected. We observed no difference in saline and sham control experiments; as such, data from the two types of controls were combined.

### Analysis of behavioral data

All analyses were performed using custom scripts in MATLAB (Mathworks). We used a bounded evidence accumulation model to fit the monkeys’ choice and RT data. The model posits that momentary evidence acquired from the stimulus is sampled sequentially and integrated until a positive or negative decision bound (±*B*) is exceeded, at which point the decision is terminated. The momentary evidence is assumed to be Gaussian with mean

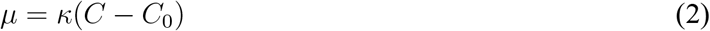

and unit variance per second, where *κ* is a constant, C is the signed motion coherence, and *C_0_* is a bias term (drift-rate offset). For the model fits in Fig. 1 and Fig. S1, we used a more parsimonious version of the model that assumes flat and symmetric decision bounds. To fit this model, we maximized the combined log-likelihood of the choice data and the mean reaction times, given the following parameters: *κ, B, C*_0_, and *t*_ND_. Where *t*_ND_ is a non-decision time parameter, which summarizes sensory and motor delays that are independent of the decision process. Details of this model and the fitting process are described in Stine et al. (2020).

We fit a more comprehensive variant of this model to test whether SC inactivation alters the decision bounds. In this variant, the decision bounds were allowed to be asymmetric and to collapse toward zero over time. The model is able to explain the full, choice-conditioned RT distributions, which, in principle, should provide a more precise estimate of the effect of SC inactivation because it takes into account all the data instead of just the mean RTs. We used the finite difference method (Chang and Cooper, 1970) to numerically solve the Fokker-Planck equation associated with the drift-diffusion process. The probability density of the accumulated evidence (*x*) as a function of time (*t*) was derived using a time-step of 0.5 ms, and we assumed that *x* = 0 at *t* = 0. The predicted decision time distribution was defined as the probability density absorbed at each bound at each time-step. This distribution was convolved with a Gaussian nondecision time distribution to compute the predicted reaction times.

The T_in_ and T_out_ decision bounds before SC inactivation were logistic functions of time:

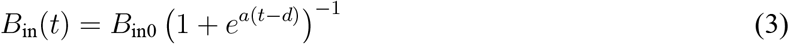

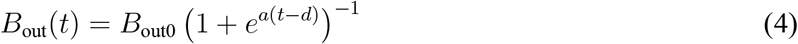

Where *a* and *d* are shape parameters and *a* is constrained to be non-negative. They are shared between the two bounds, such that the difference between the Tin bound and the Tout bound is only determined by the *B*_in0_ and *B*_out0_ parameters. A variety of other bound shapes *(e.g.* linear, hyperbolic, and exponential collapse) produced similar results.

To avoid over-fitting, we limited the number of parameters that were allowed to vary before and after SC inactivation. Such parameters are denoted by prime notation (*e.g. x*’). The first two were 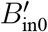 and 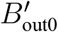, which separately scaled the corresponding decision bound after SC inactivation. The third parameter was a change in the drift-rate offset 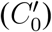, which captures any changes in choice bias that are not explained by asymmetric bounds. Finally, we allowed the non-decision time mean (*μ*) and standard deviation (*σ*) associated with T_in_ choices to increase after SC inactivation. The model has 15 parameters in total 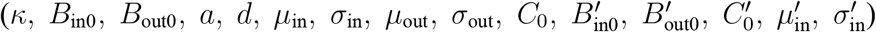. To ensure that the model was properly constrained, we empirically estimated 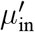 and 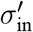 using the increase in the mean and standard deviation of saccadic latencies observed in the delayed saccade task. Recent work has shown that such constraints on the non-decision time parameters are critical to reveal cases of model misspecification (Stine et al., 2020). We fit data from before and after inactivation simultaneously, using Bayesian adaptive direct search (BADS; Acerbi and Ma, 2017) to maximize the log-likelihood of the data given the parameters.

We also summarized the effect of SC inactivation on behavior using a logistic function fit to the choice data alone. The proportion of rightward choices as a function of coherence is given by

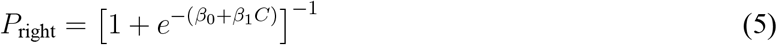

where *β*_0_ determines the left-right bias and determines the slope of the choice function. To test whether the effect of muscimol injection on bias and slope was significantly different from that of saline injection, we performed a nested model comparison. The full model was defined by the following equation:

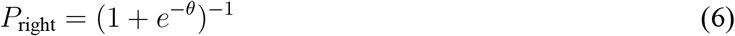

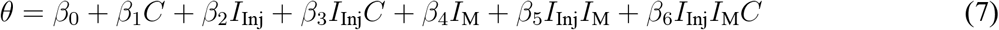

Where *I*_M_ is an indicator variable denoting the session type (saline or muscimol) and *I*_Inj_ is an indicator variable denoting whether the trial came before or after injection. We tested for a significant interaction between *I*_Inj_ and *I*_M_ (bias effect) or between *I*_Inj_, *I*_M_, and C (slope effect), by removing these terms and comparing the resulting deviance to that of the full model with a log-likelihood ratio test.

### Analysis of neural data

Single units were identified in the neural data using Kilosort 2.0 and manual curation in Phy. In LIP, we restricted our analyses to neurons with RFs that overlapped the contralateral choice target (and the SC RFs). RFs in LIP were identified post-hoc by characterizing activity in the delayed saccade task, and whether the identified RF overlapped the contralateral target was determined by visual inspection. Crucially, such determination was made before analyzing activity in the RDM task. Overall we recorded from 1,084 LIP neurons (582 from Monkey M), of which 164 (79 from Monkey M) had RFs that overlapped the contralateral choice target.

For all analyses, spike trains were convolved with a 50 ms rectangular filter. For visualization purposes, the single-trial traces displayed in Figure 3 were produced by convolving spike trains with a Gaussian kernel (σ = 25 ms). The trial-averaged responses in Figure 2A,B were computed by averaging activity over all neurons aligned to either motion onset or saccade onset. To avoid averaging over stereotyped dynamics associated with motion onset and the saccade, we implemented an attrition rule for each event-alignment and brain area. For the motion-aligned averages, we did not include spikes that occurred within 70 ms of the saccade in LIP and within 170 ms in SC. For the saccade-aligned activity, we did not include spikes that occurred within 200 ms of motion onset in both areas.

We used linear regression to analyze the effect of motion pulses on activity in each area as a function of time, relative to pulse onset. The regression includes the pulse and other factors that might affect the firing rate as a function of time on each trial.

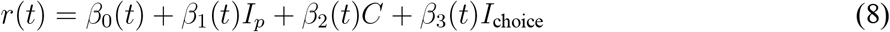

where *I_p_* is a signed indicator variable signifying the direction of the pulse (left or right), and *I*_choice_ is a signed indicator variable signifying the monkey’s choice. Fig. 2C,D depict *β*_1_(*t*), which captures the effect of the pulse on activity as a function of time. Before applying the regression, we subtracted *r*(*t*) by the average activity between 0-100 ms after pulse onset, which ensured that *β*_1_ began at a value of 0. To quantify the persistence of the pulse effect on neural activity, we fit *β*_1_(*t*) of each area with a Gaussian function with two extra parameters, which scales and shifts the function on the ordinate. The persistence is defined as the standard deviation (*σ*) of the fitted function. We tested for a significant difference in persistence between LIP and SC by fitting *β*_1_ (*t*) of both areas with a single *σ* parameter. We then compared the resulting log-likelihood to that of a model that allowed for different *σ* parameters using a likelihood ratio test.

To analyze single-trial activity, we computed the average of the simultaneously recorded neurons. These averages are standardized (z-scored), using the mean and standard deviation of activity from 0-400 ms after motion onset. The standardization ensures that any single session does not dominate subsequent analyses. Our conclusions do not depend on the specific choice of window.

We developed a simple algorithm to detect bursts of SC activity. On each trial, the algorithm first applies a threshold (3.5 σ) on the population firing rate to identify high firing-rate-events. If this first threshold is exceeded, the algorithm applies a second threshold to the derivative of the firing rate within in the epoch 80 ms before to 120 ms after the first threshold crossing. The derivative was smoothed (±50 ms running mean) before applying the second threshold (250 sp/s^2^). The event is classified as a burst if both thresholds are exceeded. The burst onset time is defined as the time when the derivative threshold is first exceeded. We hand-tuned the two thresholds and the size of the window by observing its performance on hundreds of randomly chosen trials. We used the same parameters to identify saccadic and non-saccadic bursts—the two types of bursts were differentiated only by their timing relative to the saccade (saccadic bursts were defined as those that occurred between −250 to 0 ms before the saccade). All analyses were repeated using a range of threshold values. Inevitably, doing so led to small quantitative changes in the burst frequency, onset times, and magnitudes, but the conclusions we draw are robust. Notably, the algorithm only detected the rising edge of the burst; it did not enforce that there be a subsequent decrease in activity. Thus, the burst-like activity profile that we observed in SC was not guaranteed to be discovered by the algorithm. Indeed, we applied the same algorithm, with the same parameters, to the LIP activity and observed a qualitatively different activity profile (Fig S4). Finally, we reached similar conclusions when we used more complex algorithms that also applied a threshold on the 2^nd^ derivative and/or enforced there to be a subsequent decrease in activity. Nevertheless, many aspects of the burst-detection algorithm are arbitrary—in particular, the timing of the burst onset. As such, we do not make claims about the relative timing of events in LIP and SC based on the identified onset times.

To identify upticks in LIP, we first computed the derivative of the population firing rate as a function of time, 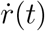, on each trial and smoothed it with a 100 ms sliding window. For the analyses in Figure 4B,C, we identified a single “uptick” on each trial, which was defined as the maximum of 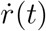 from 200 ms after motion onset to 250 ms before saccade initiation. The uptick size is the difference in firing rate 75 ms after the maximum and 75 ms before the maximum.

We used both *r*(*t*) and 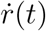 to predict, at each time-point *t*, whether the decision would terminate in a T_in_ choice 50-150 ms later. In the context of signal-detection-theory, the time-points on each trial defined as “signal present” are −50 ≤ *t* — *t*_sac_ ≤ 150 ms, where *t*_sac_ is the time of initiation of a T_in_ saccade. All other time-points are defined as having “signal absent”. The hit rate is defined as the proportion of *signal-present* time-points, across all trials, in which the quantity of interest exceeded a criterion. Likewise, the false alarm rate was defined as the proportion of *signal-absent* time-points, across all trials, in which the quantity of interest exceeded the same criterion. The quantity of interest depended on the model. It was *R*(*t*) for the firing rate model, *R*’(*t*) for the derivative model, and *β*_1_*R*(*t*—50)+*β*_2_*R*’(*t*) for the combination model. The *β* terms in the combination model were optimized using logistic regression. For each model, the quantity was compared to 100 linearly-spaced criterion values and the *d*’ was calculated using the hit rate and false alarm rate yielded by the criterion that maximized the former and minimized the latter. *d*’ was calculated as

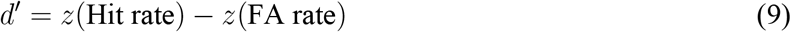

where *z* is the inverse CDF of the standard normal distribution. To calculate the change in hit rate and false alarm rate after SC inactivation (Fig 6E), we found the optimal criterion for the data before SC inactivation, and applied it to the data after SC inactivation.

## Acknowledgements

We thank Richard Krauzlis, James Herman, and Leor Katz for their invaluable advice and teachings on the SC recordings and inactivation experiments. We thank Brian Madeira, Cornel Duhaney, and Mehdi Sanayei for their help with animal care, surgery, and training, and we thank Columbia University’s ICM for the quality of care they provide for our animals, especially during the pandemic and lockdown. We would further like to thank Tanya Tabachnik and her team at the Zuckerman Institute Advanced Instrumentation Core and Tim Harris, Wei-lung Sun, Jennifer Colonell, and Bill Karsh at HHMI Janelia for their continued support with Neuropixels1.0-NHP45 probes development and testing. Finally, we thank Greg Horwitz for comments on an earlier draft of the manuscript and all members of the Shadlen lab for their feedback and engaging discussions. This research was supported by the Howard Hughes Medical Institute, an R01 grant from the NIH Brain Initiative (M.N.S., R01NS113113), a T32 and F31 grant from the National Eye Institute (G.M.S, T32 EY013933, F31 EY032791), and funding from the Grossman center (E.M.T), the Brain and Behavior Research Foundation (E.M.T and D.J.) and the Simons Foundation (D.J.).

## Author contributions

GMS and MNS designed the experiment. GMS, EMT, and DJ collected the data. GMS and MNS analyzed the data. All authors wrote the manuscript.

---

**Figure S1:**
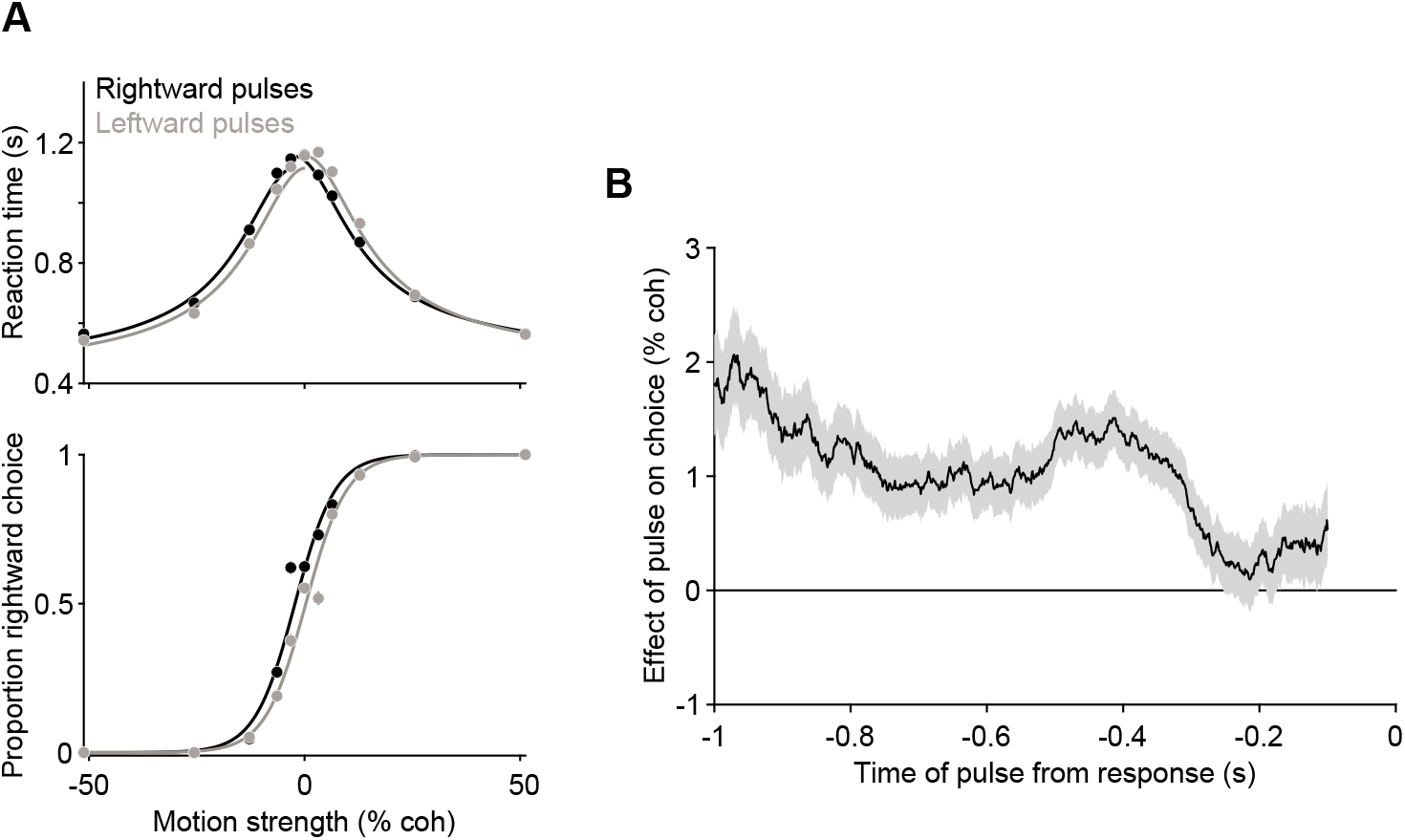
Effect of motion pulses on behavior, Related to Figure 1. **A,** Choice *(bottom)* and reaction time (*top*) data as a function of motion strength from two monkeys, plotted separately for trials with leftward (grey) and rightward (black) motion pulses. The pulses had a biasing effect equivalent to shifting the choice function left or right by ±1.4 % coh (p < .001, likelihood ratio test). **B**, Effect of motion pulses on choices as a function of time from the response. Pulses had a persistent effect on choices, consistent with temporal integration of motion evidence. Shading, standard error.

**Figure S2:**
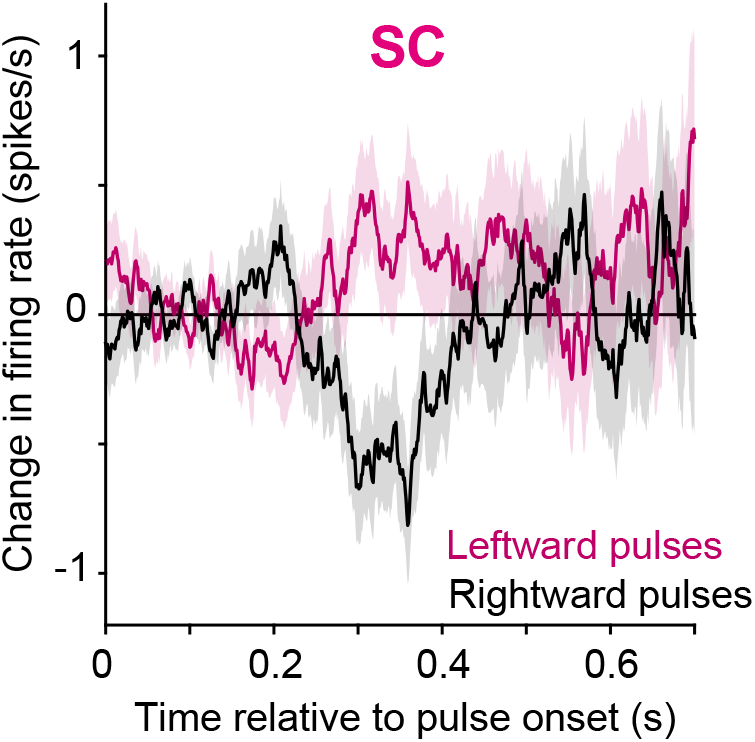
Effect of motion pulses on prelude neurons in SC, Related to Figure 2. Average activity, aligned to pulse onset, of 79 SC neurons identified as having spatially-selective persistent activity during a memory-guided saccade task (*i.e.*, visuomovement prelude neurons). Activity is baseline subtracted and plotted separately for leftward (magenta) and rightward (black) pulses. These neurons comprise a subset of those displayed in Fig. 2. (Shading shows s.e)

**Figure S3:**
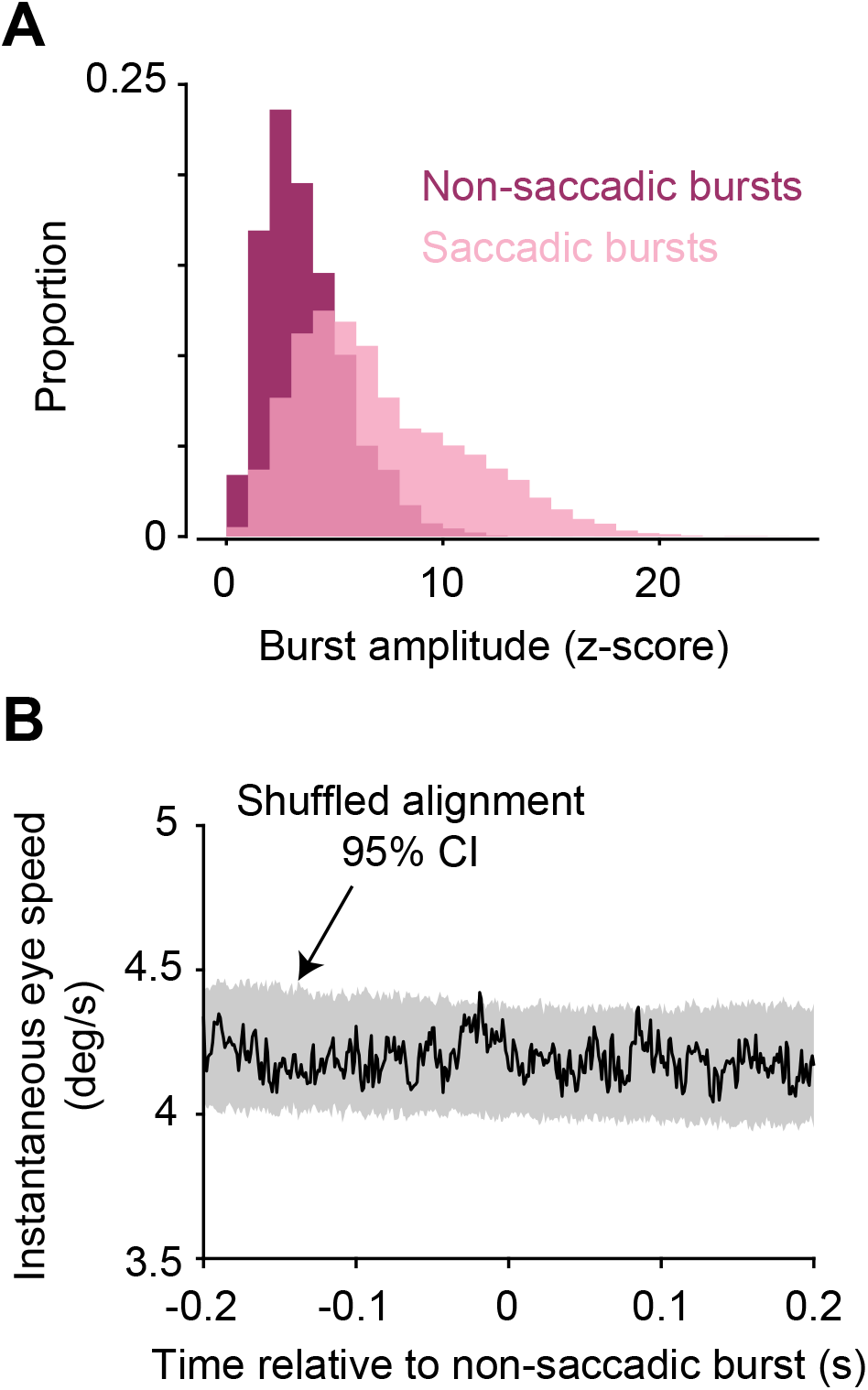
Additional analyses of SC bursts, Related to Figure 3. **A,** Distribution of burst amplitudes (peak z-score) for non-saccadic (magenta) and saccadic bursts (pink). **B**, Non-saccadic bursts are not associated with eye movements. Average instantaneous eye speed 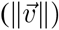 is aligned to the onset of non-saccadic bursts (black line). The shaded region shows the 95% confidence interval of the average eye speed associated with 2,000 bootstrapped, random alignments. For each trial, we chose a time-point for alignment by randomly sampling the distribution of burst-onset times, computed the average eye speed aligned to these random samples, and repeated the exercise 2,000 times.

**Figure S4:**
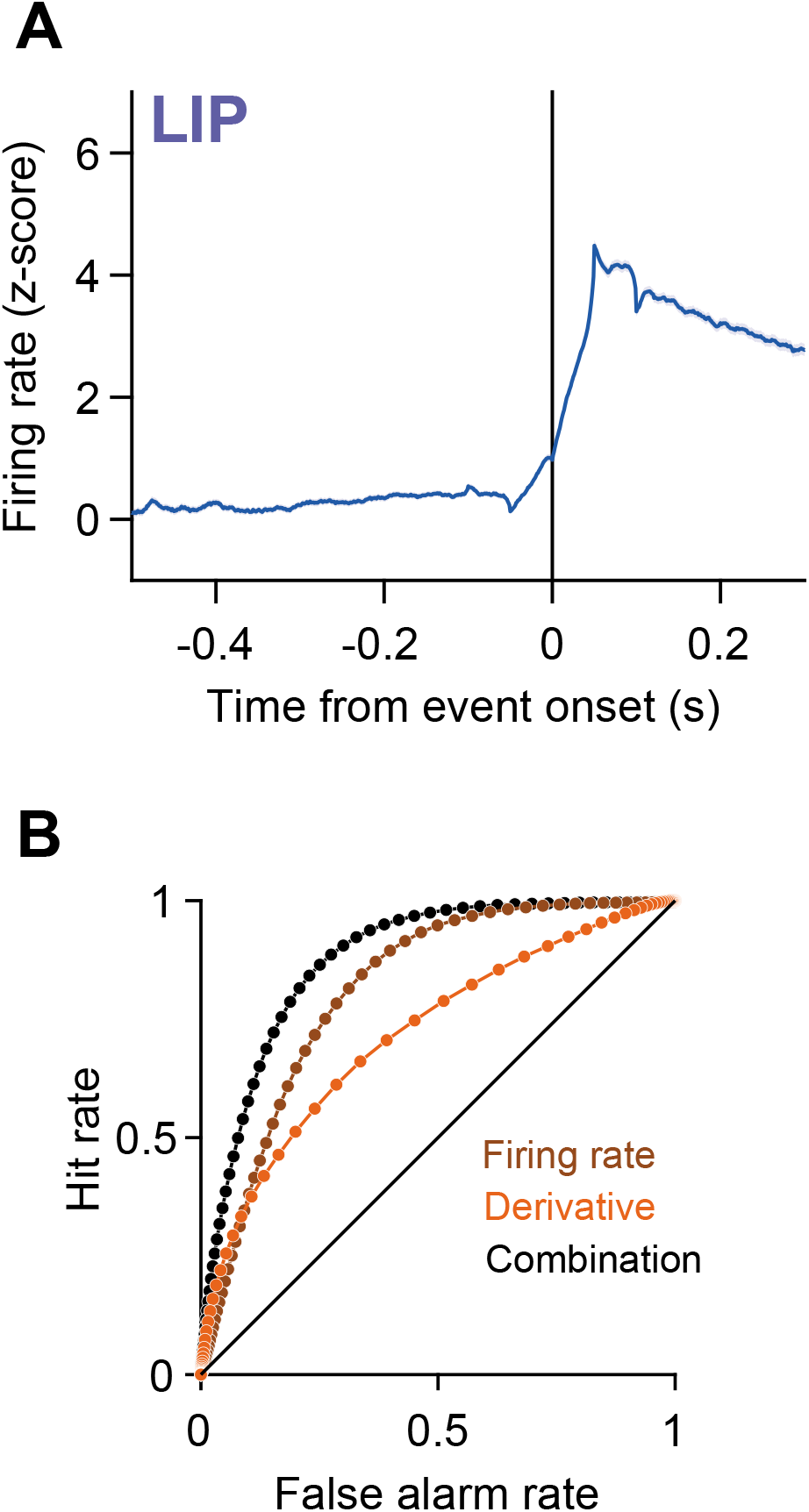
Lack of bursting in LIP, and, ROC curves associated with predictions of decision termination, Related to Figure 4. **A,** LIP activity aligned to events identified by the burst-detection algorithm. The same algorithm used to identify SC bursts in Figure 4A is applied to LIP activity here. Alignment of LIP activity to the detected events renders a qualitatively different activity profile as that observed in SC. **B**, Receiver operating characteristic (ROC) curves associated with the d’ values in Figure 4D. The hit rate and false alarm rate yielded by each model is plotted for 100, linearly-spaced criterion values. d’ values are computed using the optimal criterion (see Methods).

**Figure S5:**
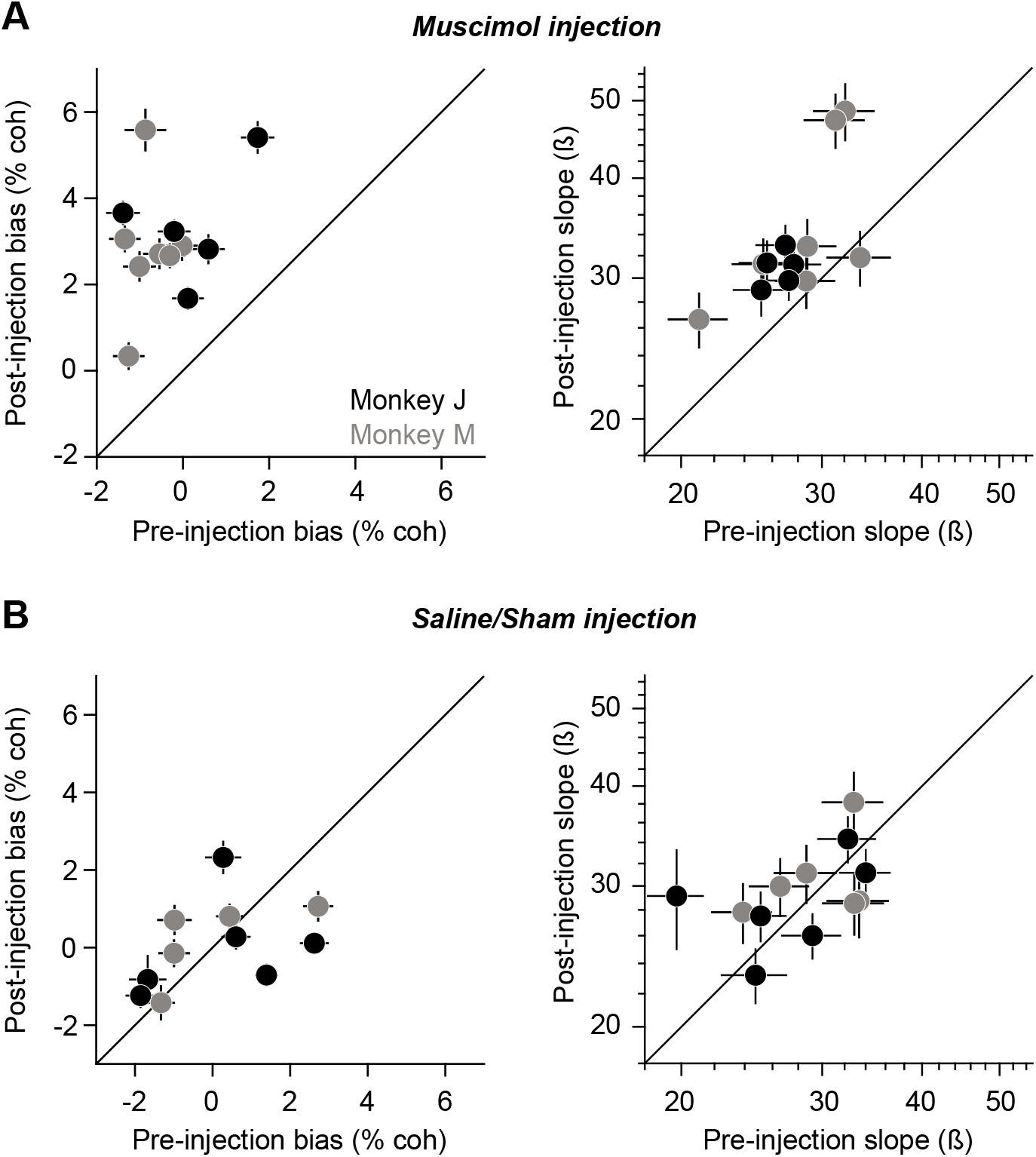
Effect of muscimol and saline injection on choice data for individual sessions, Related to Figure 5. **A**, Change in bias *(left)* and slope *(right)* following muscimol injection. Each data point depicts a single session. **B**, Same as **A** but for saline or sham injection sessions. Error bars represent standard error.

**Figure S6:**
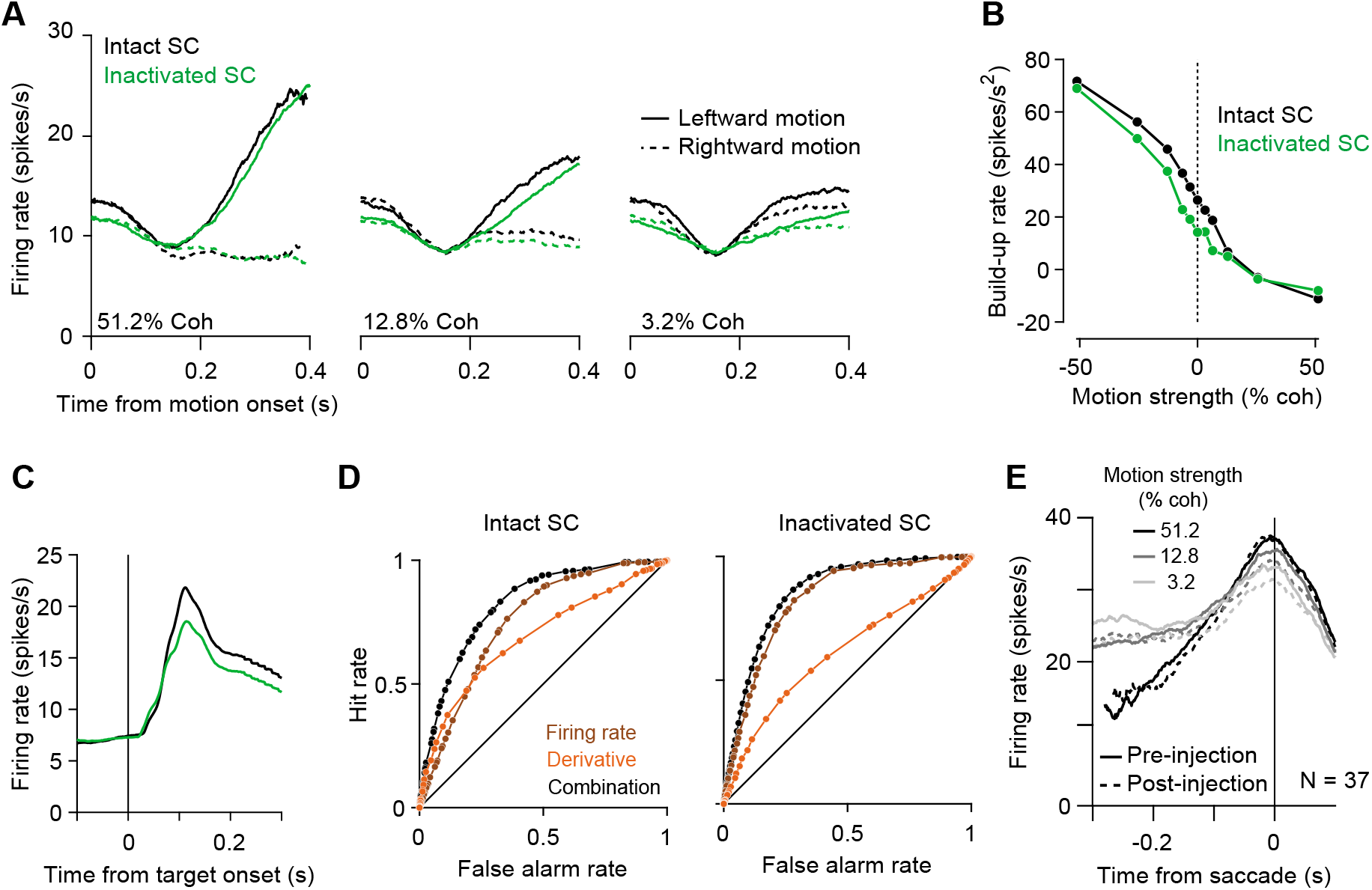
Additional effects of SC inactivation on LIP activity, and, saline control data, Related to Figure 6. **A,** Effect of SC inactivation on LIP activity early in the trial. Average of 58 LIP neurons before (black) and after (green) SC inactivation, aligned to motion onset. Activity is split by motion direction (line style) and plotted separately for a strong *(left),* intermediate *(middle),* and weak *right)* motion strength. **B**, Build-up rate of LIP activity as a function of motion strength. Negative motion strengths represent leftward motion. Build-up rates are calculated using the epoch 200 ms after motion onset to 400 ms after motion onset. SC inactivation caused a slight, overall decrease in build-up rates. **C**, Effect of SC inactivation on transient visual responses in LIP. Average LIP activity before (black) and after (green) SC inactivation is aligned to the onset of the visual targets, which occurs before the RDM stimulus is presented. **D**, ROC curves associated with the analysis in Fig. 6C. **E**, Same as Fig. 6A but for saline/sham sessions. Saline injection had a negligible effect on LIP activity at the end of the decision.

